# FK506 binding protein 5 regulates cell quiescence-proliferation decision in zebrafish epithelium

**DOI:** 10.1101/2023.03.02.530846

**Authors:** Yingxiang Li, Chengdong Liu, Xuanxuan Bai, Mingyu Li, Cunming Duan

## Abstract

The cell proliferation-quiescence decision plays fundamental roles in tissue formation and regeneration, and its dysregulation can lead to human diseases. In this study, we performed transcriptomics and genetic analyses using a zebrafish model to identify pathways and genes involved in epithelial cell quiescence-proliferation regulation. In this *in vivo* model, a population of GFP-labeled epithelial cells known as ionocytes were induced to reenter the cell cycle by a physiological stress. Transcriptomics analysis identified 1168 genes up-regulated and 996 genes down-regulated in the reactivated cells. GO and KEGG pathway analyses revealed that genes involved in transcription regulation, cell cycle, Foxo signaling, and Wnt signaling pathway are enriched among the up-regulated genes, while those involved in ion transport, cell adhesion, and oxidation-reduction are enriched among the down-regulated genes. Among the top up-regulated genes is FK506 binding protein 5 (Fkbp5), a member of the conserved immunophilin family. CRISPR/Cas9-mediated Fkbp5 deletion abolished ionocyte reactivation and proliferation.

Pharmacological inhibition of Fkbp5 had similar effects. Further analyses showed that genetic deletion and inhibition of Fkbp5 impaired Akt signaling. Forced expression of a constitutively active form of Akt rescued the defects caused by Fkbp5 inhibition. These results uncover a previously unrecognized role of Fbkp5 in regulating the quiescence-proliferation decision via Akt signaling.

**Impact Statement:** Transcriptomic and genetic deletion studies unravel a new role of Fkbp5 in promoting cell reactivation via Akt signaling.

**Highlights:**

- Transcriptomic analysis reveals several molecular pathways altered during epithelial cell quiescence-proliferation transition.
- Fkbp5 is highly up-regulated in reactivated and dividing cells.
- Fkbp5 promotes epithelial cell reactivation and proliferation via Akt signaling.

## Introduction

The active cell cycle consists of four phases: G1, S, G2, and M. In addition, the G0 phase or quiescence phase exists outside of the proliferative cycle. Cells in the quiescence phase, while non-dividing and dormant, can reenter the cell cycle upon injuries or receiving appropriate signals and stimuli (Pennycook and Barr 2020). Quiescence prevents long-lived cells (such as adult stem cells) from accumulation of genomic aberrations and stress. Dysregulation of the proliferation-quiescence balance can lead to hyper- and hypo-proliferative human diseases, including cancer and fibrosis (Pennycook and Barr 2020).

The molecular mechanisms underlying the proliferation-quiescence decision have been studied extensively using cultured cells (Pennycook and Barr 2020). Based on data collected from cultured mammalian cells kept in mitogen-free media for different time periods, Pardee (1974) proposed that a “restriction” point exists in late G1 phase (Pardee 1974). Cells that have crossed this check point prior to the mitogen removal usually complete the next cell cycle. In contrast, cells that have not reached this check point at the time of mitogen removal become quiescent and enter the G0 phase. Yao et al. (2008) showed that Rb and E2F play a critical role in regulating the proliferation-quiescence decision in cultured mammalian cells (Yao, Lee et al. 2008). Subsequent studies have led to the proposal of a bifurcation mechanism controlled by CDK2 and its inhibitor p21 (Spencer, Cappell et al. 2013). Cells that cross this bifurcation point at the end of mitosis with high CDK2 activity enter the next active cell cycle. Those with low CDK2 activity enter the quiescence phase (Spencer, Cappell et al. 2013).

These *in vitro* findings have been integrated into a two-step model recently (Pennycook and Barr 2020). According to this model, the proliferation-quiescence decision is regulated by two bistable switches. The Rb-E2F acts as the first switch. This bistable switch integrates extracellular mitogenic signals via Cyclin D:CDK4/6 activity. The Rb-E2F switch activity is also influenced by DNA damage and metabolic signals. A second bistable switch is regulated by CDK2 activity and p21 degradation. The time between these two points represents a window for cell cycle reversibility (Pennycook and Barr 2020). While these findings have provided important insights into how cells exit the active cell cycle and enter the quiescence, key questions remain. To date, the molecular nature and precise position of the commitment point in the cell cycle is unknown (Pennycook and Barr 2020). We have a limited understanding of how quiescent cells are reactivated and re-enter the active cell cycle. Moreover, these models are developed primarily based on cell culture systems, mostly in immortalized cell lines. We now understand that there are tremendous complexities *in vivo*, ranging from crosstalk with various hormones and growth factors, the availability of local nutrients and oxygen, to various feedback loops. These complexities cannot be captured by cell culture based assays.

We have recently discovered that a population of epithelial cells in zebrafish, known as Na^+^-K^+^-ATPase-rich ionocytes or NaR cells, are reactivated and undergo robust cell division in response to low calcium stress or genetic manipulations (Dai, Bai et al. 2014, Liu, Dai et al. 2017, Xin, Malick et al. 2019, Liu, Li et al. 2020; Li et al., 2021). NaR cells are structurally and functionally similar to human intestinal and renal epithelial cells and contain all molecular components for transcellular Ca^2+^ transport, including the epithelial Ca^2+^ channel Trpv6. Zebrafish live in hypoosmotic aquatic habitats and use NaR cells to take up Ca^2+^ from the surrounding water to maintain their body calcium homeostasis (Hwang 2009). Under normal medium, the constitutively open Trpv6 mediates Ca^2+^ influx continuously and maintains a high level of cytoplasmic free Ca^2+^ ([Ca^2+^]_c_) in these cells. The high [Ca^2+^]_c_ suppresses Akt-Tor signaling via the conserved protein phosphatase PP2A and promotes cell quiescence (Xin, Malick et al. 2019). When fish are transferred to an embryo medium containing low [Ca^2+^] or when Trpv6 is genetically deleted, the [Ca^2+^]_c_ drops and NaR cells re-enter the cell cycle due to the re-activation of Igfbp5a-Igf1r-PI3 kinase-Akt-Tor signaling pathway (Dai, Bai et al. 2014, Liu, Dai et al. 2017, Xin, Malick et al. 2019, Xin, Guan et al. 2021; Li et al., 2021). These findings are consistent with the known role of the AKT-mTOR signaling in reactivating adult stem cells and T cells in mammals and *Drosophila* (Chen, Wu et al. 2002, Meng, Frank et al. 2018, Kim and Guan 2019), suggesting this is an evolutionarily conserved mechanism(s).

To identify pathways and molecules regulating the quiescence-proliferation transition in *vivo*, we engineered a stable zebrafish transgenic line Tg(*igfbp5a*:GFP), in which NaR cells are genetically labeled with GFP. These transgenic fish faithfully report NaR cell reactivation (Liu, Dai et al. 2017, Xin, Malick et al. 2019, Liu, Li et al. 2020). Zebrafish are oviparous vertebrates. Their free-living, tiny, and transparent embryos make it a popular model organism in developmental biology. NaR cells in particular are located on the surface of yolk sac skin larval fish, making Tg(*igfbp5a*:GFP) fish well suited for in *vivo* analysis of the quiescence-proliferation regulation. RNA-seq analysis results showed that genes involved in cell cycle, gene transcription, Foxo signaling etc. are up-regulated in reactivated NaR cells, while genes involved in ion transport, cell adhesion, and oxidation-reduction are down-regulated. Further data mining and genetic experiments uncovered Fkbp5 as a key regulator of NaR cell quiescence-proliferation decision *in vivo*.

## Results

### The transcriptomic profile of reactivated NaR cells

The RNA-seq experiment design is shown in Figure 1A. NaR cells were isolated by FACS sorting from Tg(*igfbp5a*:GFP) fish treated with the induction medium or control medium. RNA was isolated from these cells and subjected to RNA-seq analysis. As reported previously (Liu et al., 2017), NaR cells in the induction medium treated fish underwent proliferation and formed clusters of newly divided cells in the yolk sac region, while those in the control medium remained undivided (Figure 1B). A total of 27,107 genes were detected. The genes mapping ratios were 83.5% ∼ 88.1% (Supplementary Figure 1A-1B). Differential gene expression (DEG) analysis detected a total of 2,164 DEGs. Among them, 1168 genes were upregulated and 996 were downregulated in reactivated NaR cells (Figure 1C; Supplementary Table S1).

**Fig. 1.**
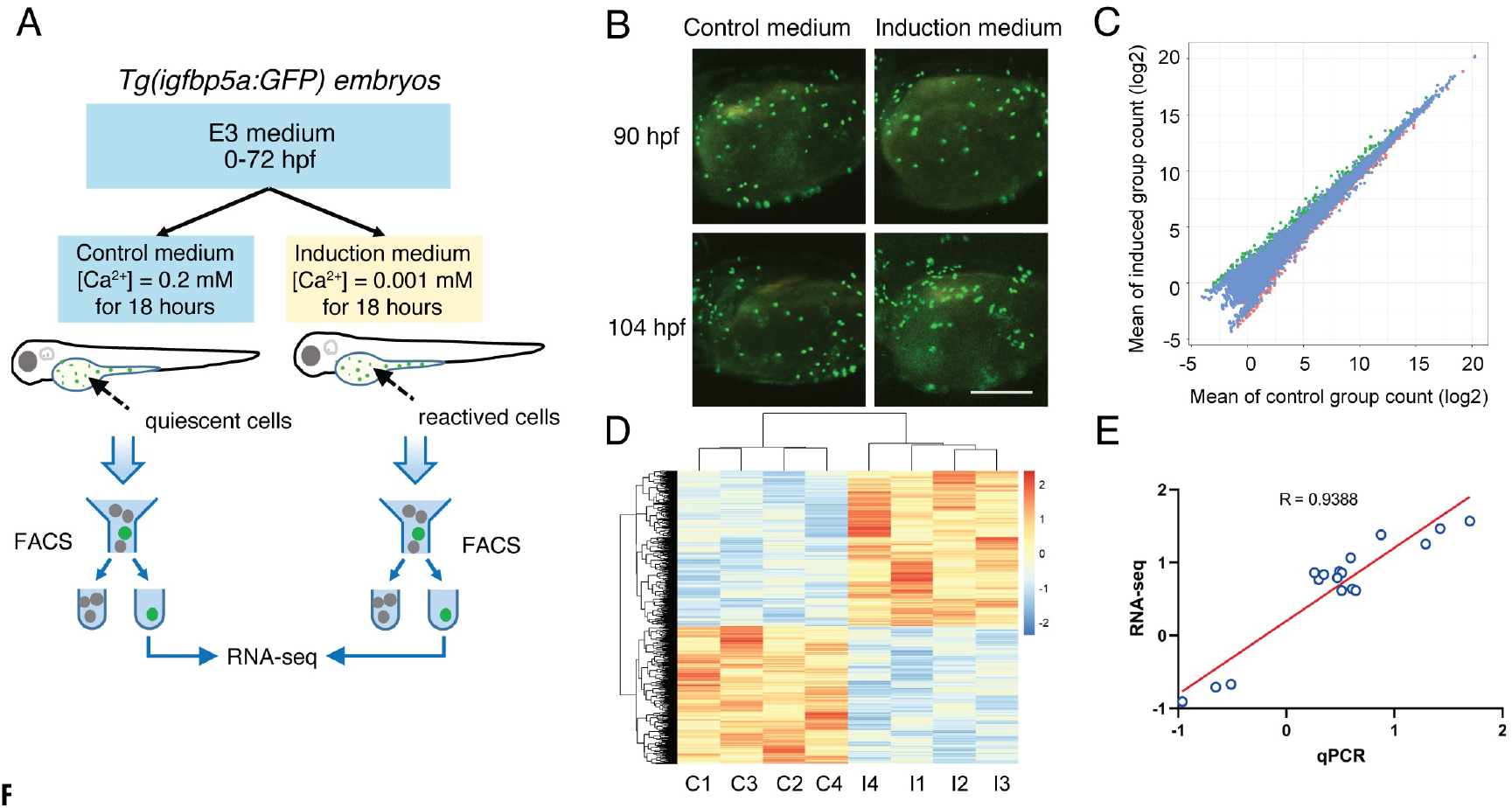
Identification of differentially expressed genes (DEGs). (A) The RNA-seq experiment design. (B) Representative images of Tg(*igfbp5a*:GFP) larvae at the indicated stages grown in the control medium or the induction medium. Shown here and in following figures are lateral views of the yolk sac region. Anterior to the left and dorsal up. Note clusters of newly divided NaR cells induced by the induction medium at 104 hours post fertilization (hpf). (C) Pairwise comparison of gene expression abundance between the control and the induced NaR cells. The up- and down-regulated genes were shown in green and red, respectively. Genes that were not changed are shown in blue. (D) Hierarchical clustering of the differentially expressed genes (DEGs) in the four control (C) and four induction groups (I) of NaR cells. Red color indicates up-regulated genes and blue color down-regulated genes. (E) qRT-PCR confirmation of RNA-seq data. Changes (log2) in the mRNA levels of the indicated genes measured by RNA-seq were plotted against those detected by qPCR. The line indicates the linear correlation between the results of RNA-seq and qPCR.

Hierarchical clustering of DEGs showed that the reactivated NaR cell datasets clustered distinctly from the control NaR cell dataset (Figure 1D). In agreement, Principal component analysis (PCA) showed that the reactivated NaR cell datasets clustered distinctly from those of the quiescent NaR cell dataset (Supplementary Figure 1C-1D). To confirm the RNA-seq results, qRT-PCR assays were performed using a different set of FACS-sorted NaR cells and similar changes were detected in 17 out of the 17 genes examined (Figure 1E; Supplementary Figure S2).

### Gene ontology (GO) and Kyoto Encyclopedia of Genes and Genomes (KEGG) enrichment analysis

The identified DEGs were annotated with GO terms and sorted based on functional categories. Genes involved in transcription regulation, oxidation reduction, multicellular organism development, DNA repair, and cell proliferation are enriched (Figure 2A). When the up-regulated and down-regulated DEGs were analyzed separately, genes involved in transcription/regulation, multicellular organism development, cell cycle/cell division, DNA repair, and Wnt signaling are enriched in the up-regulated DEGs (Figure 2B). Among the down-regulated genes, the most enriched GO term is transport, followed by oxidation reduction, cell adhesion, and vesicle-mediated transport (Figure 2C).

**Fig. 2.**
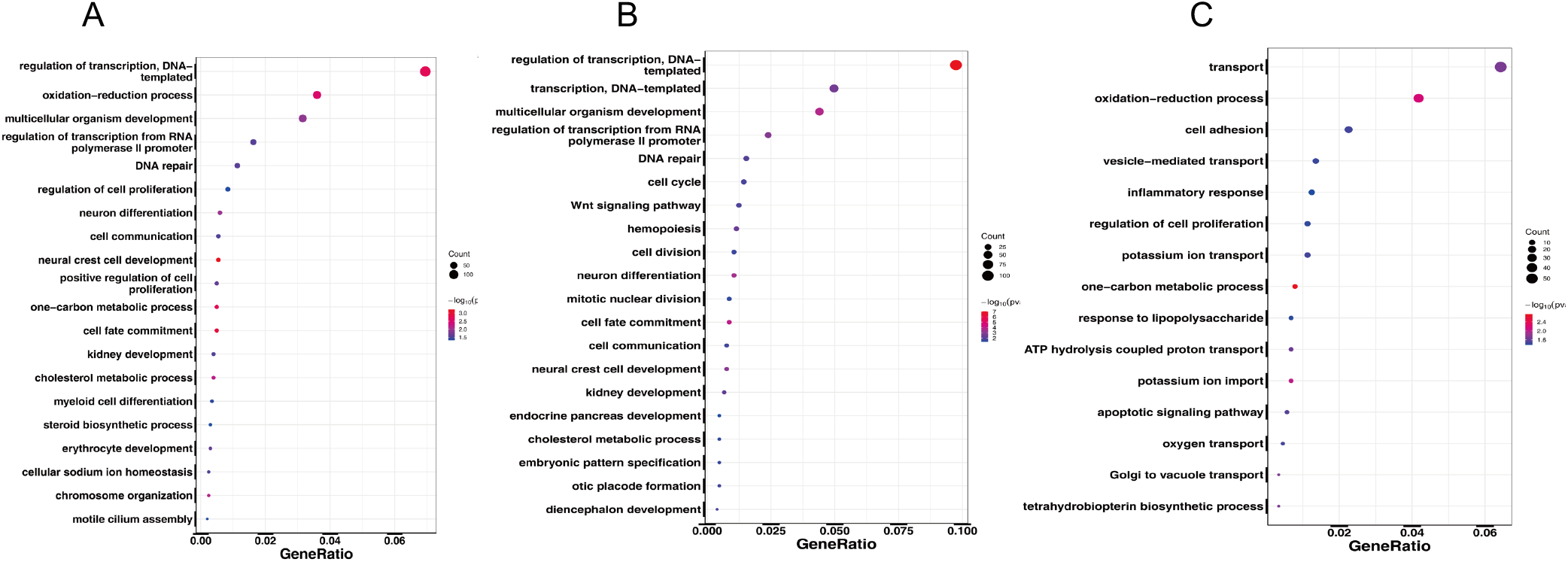
GO enrichment analysis. GO analysis results of all (A), up-regulated (B), and down-regulated (C) DEGs. The dot size indicates the number of genes, and the x-axis indicates the identified genes ratio in the relevant GO term.

KEGG pathway enrichment analysis was performed next. The DEGs were significantly enriched in cell cycle, Foxo signaling, p53 signaling, PPAR signaling, and Fanconi anemia pathway (Fig. 3A). These pathways were also enriched in the up-regulated DEGs (Figure 3B). Additionally, Wnt signaling, cellular senescence, and melanogenesis genes were also enriched (Figure 3B). The enrichment of upregulated “cell cycle” genes is in good agreement with the fact that these are rapidly dividing cells. Although KEGG analysis suggested an enrichment of the p53 signaling and cell senescence pathways (Figure 3A-3B), many of the genes are overlapping with those in the cell cycle pathway (Supplemental Fig. S3). Genes in this category include *cdk21, gadd45aa, ccne2, cdkn1a,ccnb2, ccnd1, chek1, e2f2* etc. (Supplementary Fig. S3). There are also significant overlaps in the Wnt and melanogenesis pathways (Supplemental Fig. S5), including *wnt7ba, wnt8b, wnt 11, wnt6b, wnt7bb* and *wnt7aa* etc., consistent with the role of Wnt signaling in regulating melanocyte development (Vibert, Aquino et al. 2017).

**Fig. 3.**
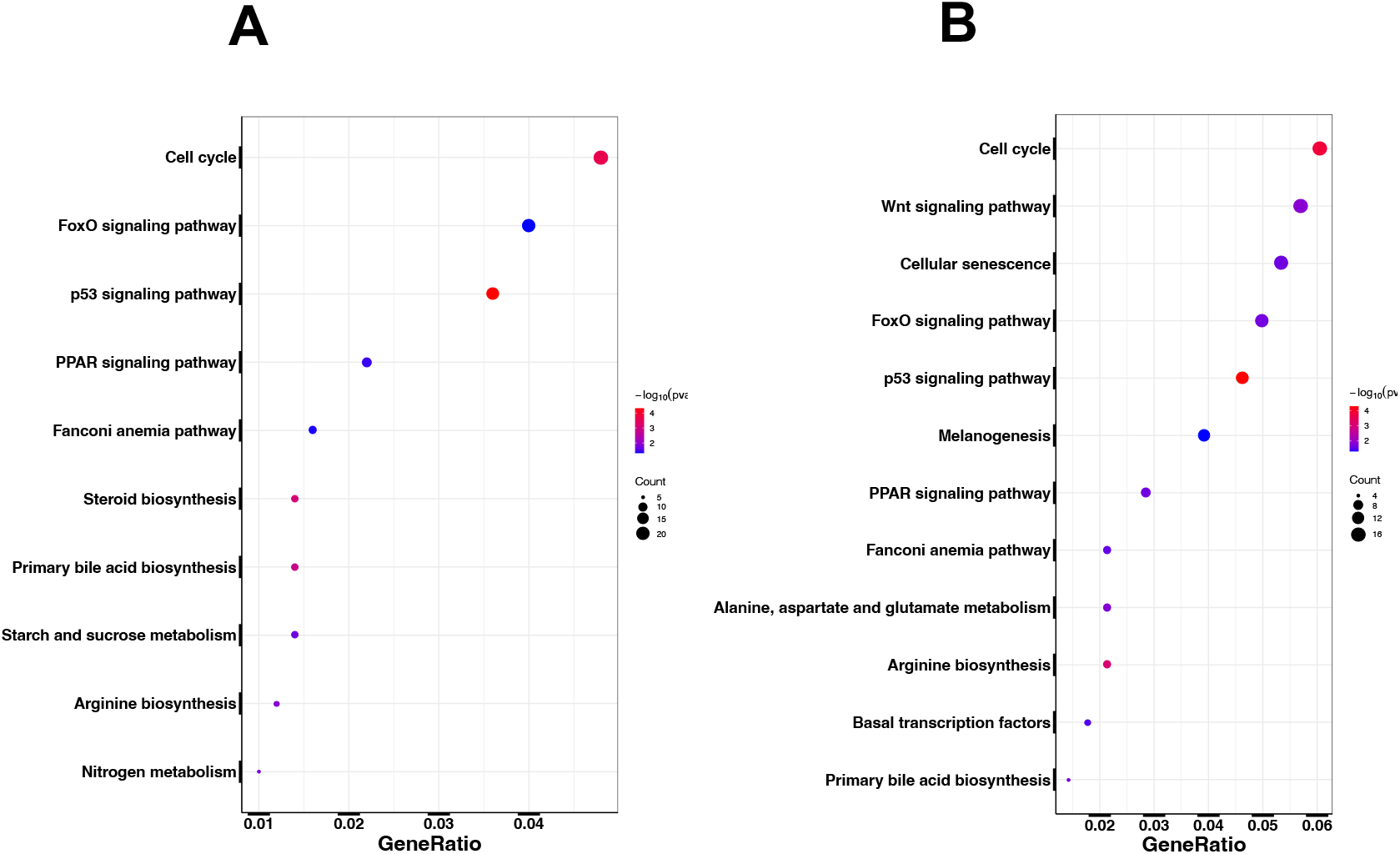
KEGG enrichment analysis. KEGG analysis of all (A), and up-regulated (B). The dot size indicates the number of identified genes, and the x-axis indicates the genes ratio in the relevant KEGG pathway.

### Fkbp5 is up-regulated and is indispensable in NaR cell reactivation

We mined the RNA-seq dataset further by ranking the upregulated DEGs based on their relative mRNA abundance in NaR cells (Table 1). As expected, the calcium channel gene *trpv6*, which has been shown to regulate NaR cell proliferation-quiescence decision by suppressing the Igf1r-PI3 kinase-Akt-Tor signaling pathway (Xin, Malick et al. 2019), is among this group. Others are *col2a1a, caspb, her9, ankhb, pck1, fkbp5, si:dkey-22i16*.*3, ponzr3*, and *wu:fj16a03* (Table 1). We were particularly intrigued by Fkbp5, a member of the conserved FK506 binding protein (FKBP) family. The FKBP family consists of a large number of structurally related proteins known for their capability of binding to and modulating the immunosuppressive actions of FK506 as well as rapamycin, a well-known mTOR inhibitor (Tong and Jiang 2016). Rapamycin treatment inhibited NaR cell reactivation and proliferation (Liu, Dai et al. 2017). Among the 19 zebrafish *fkbp* gene family members (Figure 4A), the expression of *fkbp5* and *fkbp-like*, but not other *fkbp* genes, were increased in reactivated NaR cells (Figure 4B). The upregulation of *fkbp5* mRNA expression was confirmed by qRT-PCR (Figure 4C). In comparison, the mRNA levels of *fkbp7*, a related member of the Fkbp gene family, showed no change (Figure 4C).

**Table 1.**
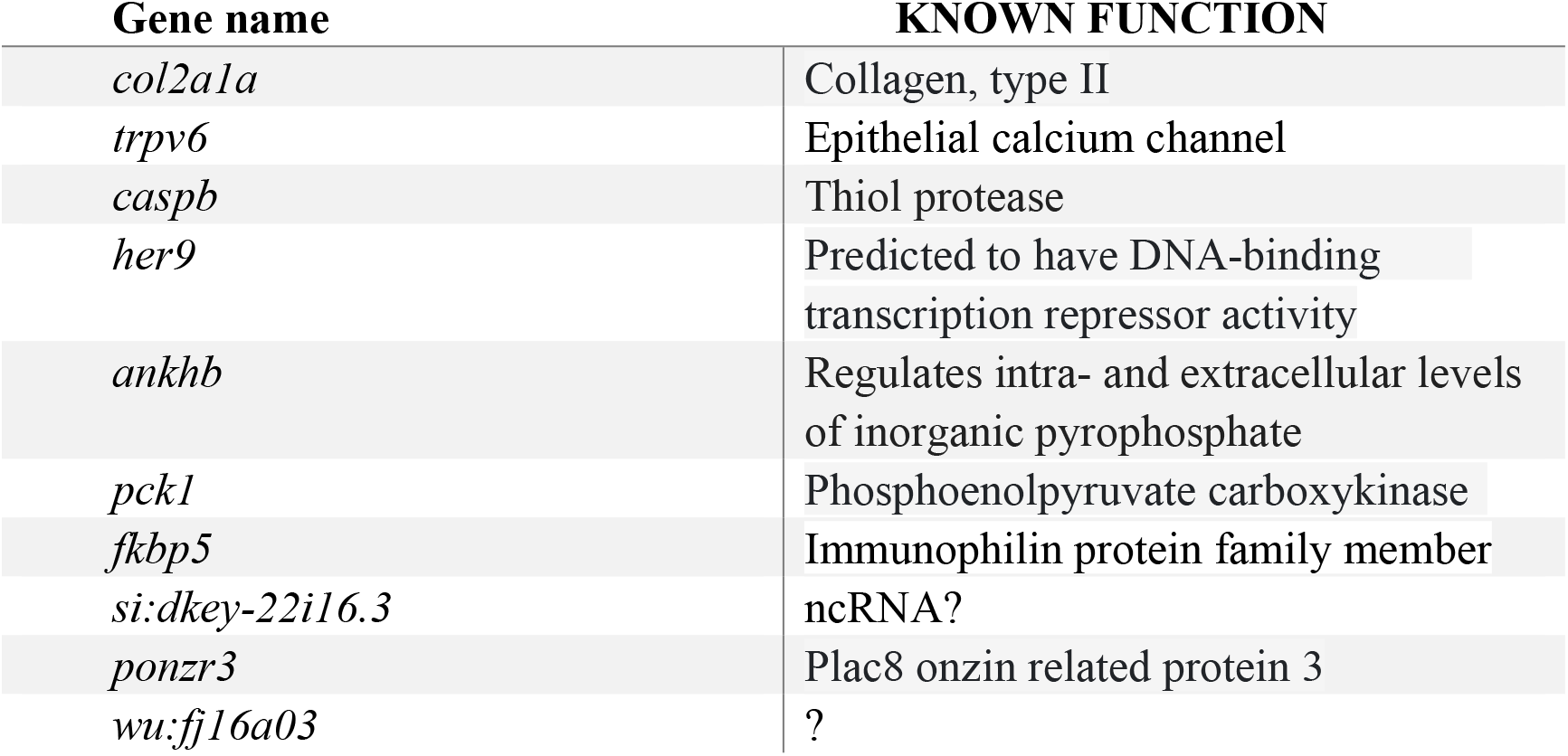
Top up-regulated genes based on mRNA abundance

**Fig. 4.**
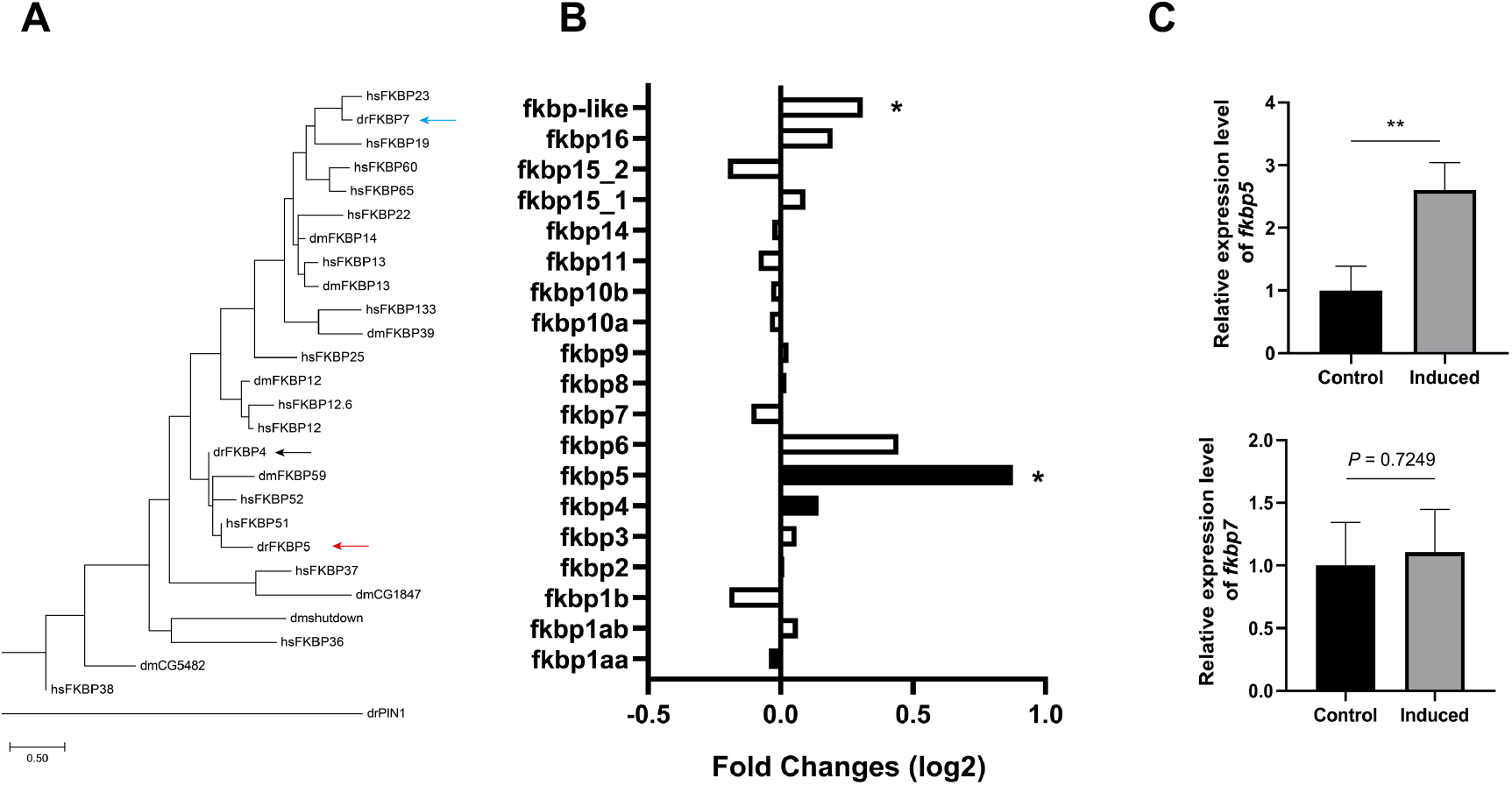
Up-regulation of fkbp5 expression in reactivated NaR cells. (A) Phylogenetic tree of humans (hs), zebrafish (dr), and Drosophila (dm) FKBPs. Pin1 prolyl *cis–trans* isomerase drPIN1is used as an outgroup. (B) Relative levels of *fkbp* mRNAs detected by RNA-seq. The mRNA levels of the indicated genes in reactivated NaR cells were normalized by those of the control NaR cells. Data shown are mean. * *P* < 0.05. (C) qRT-PCR results. NaR cells were isolated by FACS as described in Fig.1. The mRNA level of *fkbp5* (upper panel) and *fkbp7* (lower panel) were detected by qRT-PCR and expressed as fold change over the control group. n=4. Data shown are mean ± SD. ** indicates *P* < 0.01.

To determine whether Fkbp5 plays a role in reactivating NaR cells, *Tg(igfbp5a:GFP)* fish were treated with FK506, a pan FKBP inhibitor (Yeh, Bierer et al. 1995). FK506 treatment inhibited the induction medium-induced NaR cell proliferation, whereas it did not change the basal NaR cell number in fish kept in the control medium (Figure 5A and 5B). Next, a CRISPR-Cas9-based F0 gene deletion approach was used to genetically delete Fkbp5. This method can convert over 90% of injected embryos directly into biallelic knockout (Wu, Lam et al. 2018, Kroll, Powell et al. 2021) and is well suited for functional analysis of essential genes. Injection of *fkbp5* targeting gRNAs abolished NaR cell reactivation, while scrambled gRNAs had not such an effect (Figure 5C). Deletion of Fkbp5 did not alter the number of NaR cells under the control medium (Figure 5C). In comparison, CRISPR/Cas9-mediated knockdown of Fkbp7 had little effect (Figure 5D). Together, these data suggest that elevated Fkbp5 expression promotes NaR cell quiescence-proliferation transition.

**Fig. 5.**
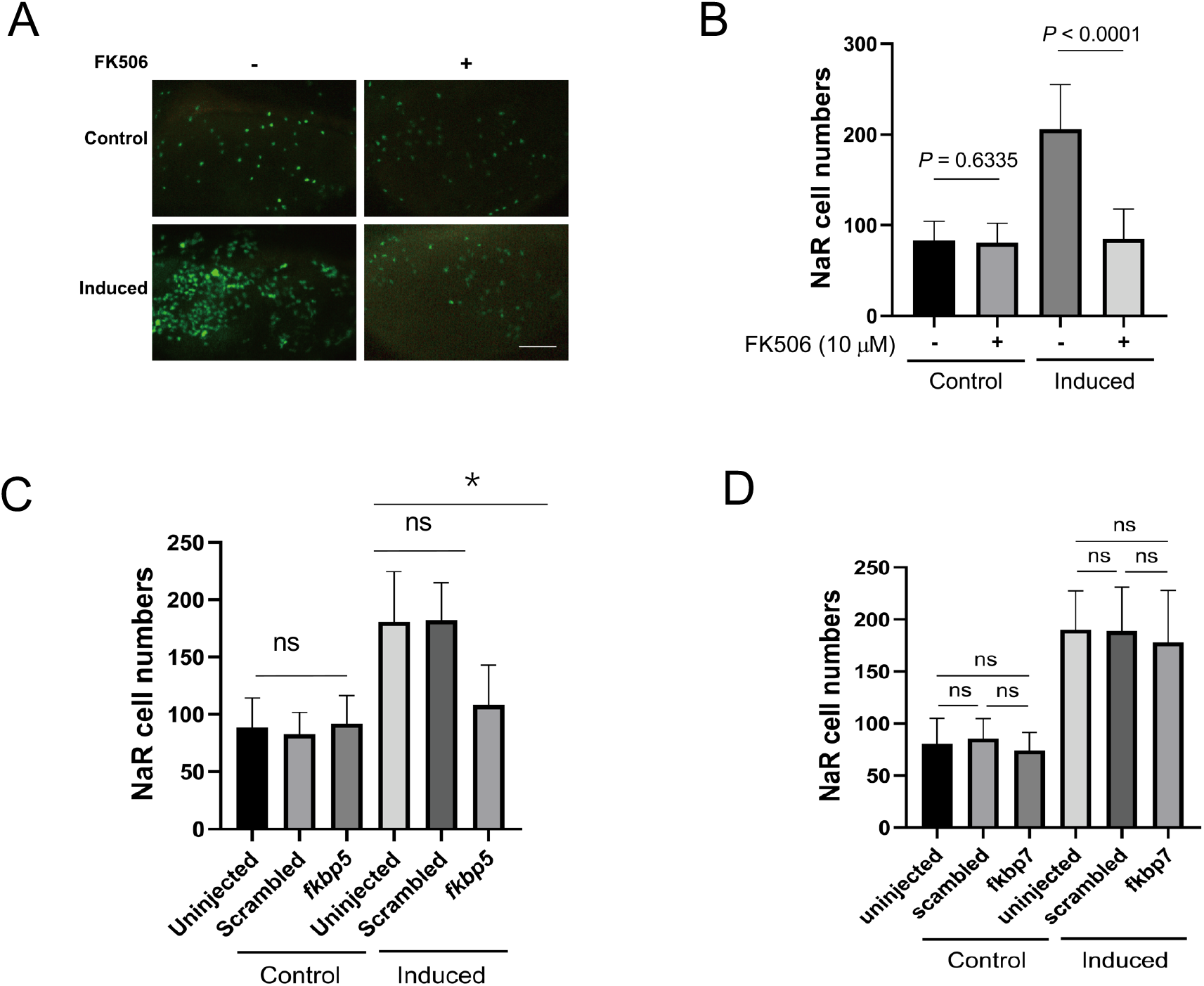
Fkbp5 promotes NaR cell quiescence-proliferation transition. (A-B) Effect of FK506. *Tg(igfbp5a:GFP)* embryos were raised in E3 medium to 3 dpf and transferred to the control and induction medium with or without 10 μM FK506. NaR cells were quantified at 5 dpf. Representative images were shown in (A) and quantified data in (B). Scale bar = 0.2 mm. Data shown are mean ± SD. n = 37∼58. (C-D) *Tg(igfbp5a:GFP)* embryos were injected with *Cas9* mRNA and gRNAs targeting *fkbp5* (C), *fkbp7* (D) or scrambled gRNA at the one-cell stage. They were raised in E3 solution to 3 dpf and transferred to the control (Control) or induction medium (induced). NaR cells were quantified 2 days later. Data shown are mean ± SD. n = 17∼69.

### Fkbp5 promotes NaR cell proliferation via Akt signaling

If Fkbp5 acts in the Igf1r-PI3 kinase-Akt pathway to stimulate NaR cell reactivation and proliferation, then deletion of Fkbp5 should alter Akt signaling activity in these cells. Likewise, constitutive activation of Akt should reverse the effects of Fkbp5 inhibition. Indeed, while the induction medium treatment resulted in a robust increase in the number of phosphor-Akt positive cells in un-injected or scrambled gRNA injected fish (Figure 6A), this effect was impaired by CRISPR/Cas9-mediated *fkbp5* deletion (Figure 6B). Next, myrAkt, a constitutively active form of Akt (Kohn, Takeuchi et al. 1996), was expressed in a subset of NaR cells using a Tol2 transposon-mediated genetic mosaic assay (Liu, Xin et al. 2018).

**Fig. 6.**
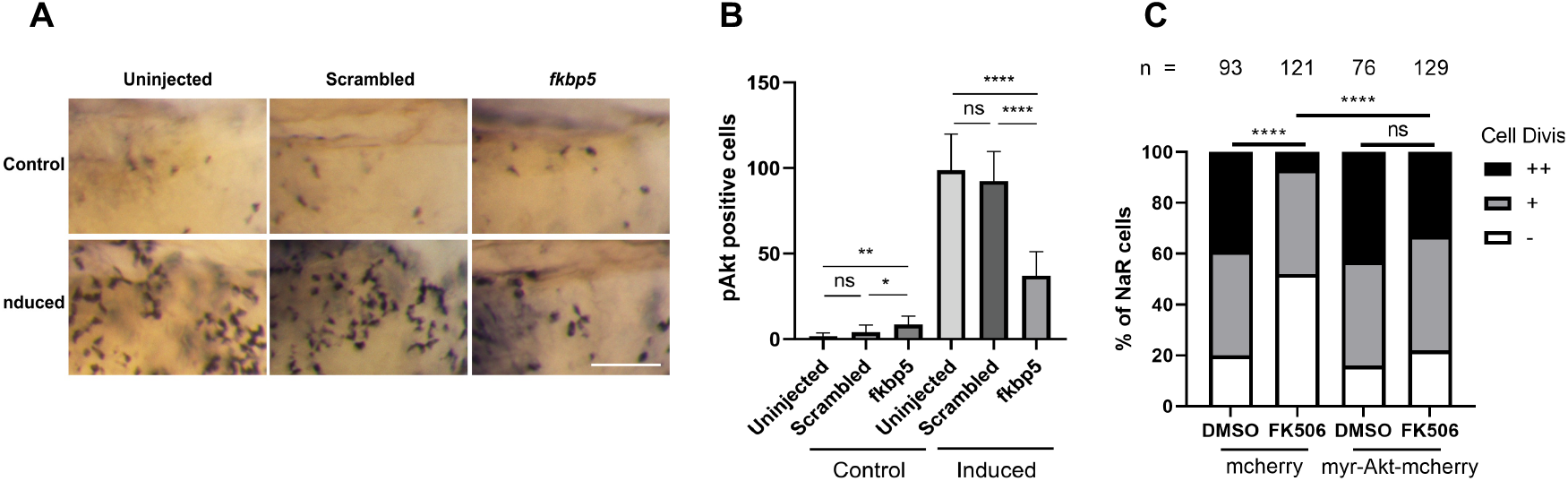
Fkbp5 promotes NaR cell quiescence-proliferation transition via Akt signaling. (A-B) *Tg(igfbp5a:GFP)* embryos were raised in E3 medium until 3 dpf and transferred to the control (Control) or induction medium (induced). Two days later, they were subjected to phospho-Akt immunostaining. Phospho-Akt-positive cells in the yolk sac region were quantified. Representative images were shown in (A) and quantified data in (B). Data shown are mean ± SD. (C) Effect of myrAkt expression. *Tg(igfbp5a:GFP)* embryos injected with *BAC (igfbp5a:mCherry)* or *BAC (igfbp5a:myr-Akt-mCherry)* were raised. At 3 dpf, they were transferred to the induction medium containing DMSO or 10 mM FK506. At 5 dpf, NaR cells labeled by both GFP and mCherry were scored following a previously established scoring system (Liu, Xin et al. 2018). NaR cells that that did not divide, divided one time or two times are scored as −,+and ++, respectively. \**P*<0.05; \*\*\*\**P*<0.0001; ns, not significant. Total cell number is shown above each column.

While FK506 treatment impaired NaR cell reactivation and proliferation in normal NaR cells, this effect was reversed by myrAkt expressing (Figure 6C). These data suggest that Fkbp5 promotes NaR cell reactivation and proliferation via Akt signaling.

## Discussion

In this study, we used a zebrafish model and mapped transcriptomic changes during epithelial cell quiescence to proliferation transition *in vivo*. Our transcriptomic analysis showed that genes involved in transcription/regulation, cell cycle, Foxo, Wnt, and PPAR signaling are increased, while those involved in ion transport, cell adhesion, and oxidation-reduction are down-regulated in the reactivated cells. These findings support the notion that low calcium stress induces differentiated NaR cells to re-enter the active cell cycle via reactivating the PI3 kinase-Akt pathway (Xin et al., 2019; 2021; Liu et al., 2020). Further data mining, genetic, and biochemical analyses showed that Fkbp5 plays a critical role in stimulating NaR cell proliferation-quiescence transition via Akt signaling.

As expected, a significant enrichment of the cell cycle genes was observed in reactivated and dividing NaR cells. Although p53 signaling and cellular senescence pathway are also enriched, a close look suggested that many of the enriched genes are overlapping with those in the cell cycle genes (Supplemental Figure S). GADD45 and CDK1 have been implicated in the modulation of cell cycle and apoptosis in mammalian cells (Kleinsimon, Longmuss et al. 2018). Likewise, ccne2, *ccnd1*, and ccnd2, which encode Cyclin E2, D1, and B2, are key regulators of cell cycle and frequently overexpressed in cancer cells (Joo, Kang et al. 2001, Nagasawa, Onda et al. 2001, Fu, Wang et al. 2004). In zebrafish, Cdk21 has been shown to promote male germ cell proliferation and meiosis (Webster, Henke et al. 2019). In addition to cell cycle genes, several DNA repair pathway genes are enriched in the reactivated and proliferation NaR cells. Cell division requires the faithful copying of the genome once per cell cycle. Robust cell replication can lead to replication stress and activates DNA damage repair pathways such as homologous recombination (Gaillard et al., 2015). It is possible the rapid dividing NaR cells may experience replication stress. This is consistent with the enrichment of Fanconi anemia genes in these cells. The Fanconi anemia pathway functions as a DNA damage repair system (Datta and Brosh, 2019).

In previous studies, we have shown that the reactivation of Akt is required and sufficient in promoting NaR cell reactivation and proliferation (Dai, Bai et al. 2014, Liu, Dai et al. 2017, Xin, Malick et al. 2019, Liu, Li et al. 2020). In this study, many genes in the Foxo signaling pathway are enriched (Supplemental Fig. S4). A key mechanism in the regulation of FOXO proteins is the phosphorylation by AKT, in response to insulin or IGFs stimulation. This modification promotes FOXO proteins to exit from the nucleus and decreases the expression of the FOXO target genes (Manning and Cantley 2007). Among the DEGs in this pathway are Sgk1 and Sgk3, two Akt-like protein kinases. SGK1 has been reported to be activated by insulin and IGFs via the PDK1 and TORC2 (Castel, Ellis et al. 2016, Zhou, Zhang et al. 2021). SGK1 increases the protein abundance and/or activity of several ion channels, solute carriers, and Na+/K+-ATPases (Lang and Shumilina 2013). Overexpression of SGK3 in hepatocellular carcinoma has been shown to increase cell cycle progression through G1 by inactivating glycogen synthase kinase-β (GSK3-β) and stabilizing CCND1 (Hou, Lai et al. 2015). Future genetic studies are needed to elucidate the possible roles of Sgk1 and/or Sgk3 in NaR cell quiescence-proliferation regulation.

Multiple Wnt ligands are enriched in the up-regulated genes in proliferating NaR cells. Wnt signaling plays a crucial role in melanocyte development (Vibert, Aquino et al. 2017). In Xenopus epidermis, Wnt signaling is required for cilia formation and differentiation of multiciliate epithelial cells. In basal cells, Wnt signaling prevents specification of epithelial cell types via *ΔN-TP63*, a master transcription factor (Haas, Vázquez et al. 2019). NaR cells are located on the yolk sac epidermis in zebrafish embryos and larvae. It will be interesting to determine whether these Wnt ligand genes play a direct role in regulating NaR cell development and proliferation. Another pathway of interest is PPAR signaling. Several genes, including *pck1* and *pck2*, are enriched in the reactivated NaR cells. While PCK1 is well known as a gluconeogenic enzyme, recent studies show that PCK1 and its mitochondrial counterpart PCK2 also increase TCA cycle flux to stimulate cell proliferation in colon cancer-derived cell lines (Balsa-Martinez and Puigserver 2015, Montal, Dewi et al. 2015, Vincent, Sergushichev et al. 2015). It is unclear whether any of these PPAR genes plays a role in regulating NaR cell fate.

NaR cells are one of the several types of ionocytes derived from epidermal stem cells (Hwang, 2009; Xin et al., 2019). NaR cells are polarized cells with an apical opening facing the external environments and basolateral sides facing the internal body fluids. These cells are responsible for transcellular Ca^2+^ transport, which also involves Na^+^ and K^+^ trafficking (Hwang 2009). Our transcriptomic analysis showed that the GO term of “transport” was most highly enriched among the down-regulated genes. Out of the 996 down regulated genes, 57 are in this group. Within this category, 11 genes are known to be involved in ion transport. These include several calcium ion channels, potassium channels, and Na^+^, Cl^-^, and H^+^ channels and transporters. The down-regulation of these genes, together with the reduction of cell adhesion genes, supports the notation that NaR cells are dedifferentiated and reactivated. We have previously found that the epithelial calcium channel *trpv6* plays a critical role in stimulating NaR cell quiescence. The present study identified three additional calcium channels, including *cacng1a* (calcium channel, voltage-dependent, gamma subunit 1a), *cacng3a*, and *trpv1* down-regulated in reactivated NaR cells. Trpv1 is critical for heat sensing (Gau, Poon et al. 2013). Currently, little is known about the function of *cacng1a* and *cacng3a*. The roles of these channels in NaR cell quiescence-proliferation balance, if any, are unknown at present and deserves further investigation.

A key finding made in this study is that Fkbp5 played a critical role in promoting NaR cell quiescence-proliferation transition. This conclusion is supported by several lines of evidence. First, *fkbp5* mRNA expression is upregulated in reactivated NaR cells. In comparison, the expression of *fkbp7* and many other *fkbp* genes was not changed. Second, pharmacological inhibition of Fkbp prevented NaR cell quiescence-proliferation transition. Furthermore, genetic deletion of Fkbp5 impaired NaR cell reactivation and proliferation. This effect is specific because deletion of Fkbp7 had little effect. Although a previous study reported that FKBP51 (encoded by the human *FKBP5* gene) promotes the expression of epithelial-to-mesenchymal transition (EMT) genes (Romano, Staibano et al. 2013), a role of Fkbp5 in cell quiescence-proliferation decision has not been reported. FKBPs have a variety of molecular masses that are involved in several signal transduction pathways (Somarelli, Lee et al. 2008, Tong and Jiang 2016, Boonying, Joselin et al. 2019). Human FKBP51 was originally discovered as a co-chaperone of Hsp90 and a component of the Hsp90-steroid receptor heterocomplex (Smith, Sullivan et al. 1993). FKBP51 contains 2 FKBP domains and 3 tetratricopeptide-repeats (TPR) domain. While the first FKBP domain (FK1) has peptidylprolyl isomerase (PPI) activity and provides a binding pocket for FK506 and rapamycin, FK2 lacks the PPIase activity. Among the FKBP family members, FKBP51 is most closely related to FKBP52 (encoded by the human *FKBP-4* gene) (Wang, 2011). Despite the high sequence similarity, these two proteins had different and often opposite roles in several biological processes. While one was found to facilitate the dephosphorylation of Akt via the phosphatase PHLPP (Pei, Li et al. 2009, Hou and Wang 2012), the other had a capability to promote Akt phosphorylation through PI3K/PDK1 and mTORC2 (Mangé, Coyaud et al. 2019). In this study, we provided genetic evidence that Fkbp5 acts upstream of Akt to promote NaR cell reactivation and proliferation. This conclusion is supported by the fact that 1) CRISPR/Cas9-mediated deletion and pharmacological inhibition of Fkbp5 attenuated Akt signaling in NaR cells, while targeted expression of myrAkt “rescued” FK506 effect on NaR cell reactivation.

As a central node for signal transduction and proliferation, Akt integrated multiple intracellular signaling including Ca^2+^ flux. Intracellular Ca^2+^/calmodulin signaling has been implicated in Akt activation in cultured mammalian cells (Deb, Coticchia et al. 2004). Our recent study showed that ER Ca^2+^ efflux could activated Ca2+/calmodulin-dependent protein kinase kinase (CaM-KK) and this in turn activated Akt to induce NaR cell reactivation (Xin, Guan et al. 2021). Human FKBP 12 and FKBP12.6 are both known to regulate ER Ca^2+^ flux via the association with ER Ca^2+^ channels like ryanodine receptors (RyRs) and inositol 1,4,5-trisphosphate receptors (IP3Rs) (MacMillan 2013, Gonano and Jones 2017). Our transcriptomic analysis showed the mRNA levels of *fkbp1aa, fkbp1ab* and *fkbp1b*, which encode zebrafish Fkbp12 and Fkbp12.6a and -b, were not significantly changed in reactivated NaR cells.

Treatment of zebrafish larvae with ryanodine receptors (RyR) inhibitors ryanodine and dantrolene blocked NaR cell reactivation, while the IP3R inhibitor xestospongin C did not have such an effect (Xin, Guan et al. 2021). Future studies are needed to determine whether Fkbp5 regulates Akt signaling by altering RyRs-mediated ER Ca^2+^ flux in NaR cells.

## Materials and Methods

### Chemicals and Reagents

All chemical reagents were purchased from Fisher Scientific (Pittsburgh, PA) unless stated otherwise. Liberase TM was purchased from Sigma-Aldrich (St Louis, MO). FK506 was purchased from Selleck Chemicals (Houston, TX). Oligonucleotide primers, the TRIzol reagent, Moloney murine leukemia virus (M-MLV) reverse transcriptase and RNaseOUT Recombinant Ribonuclease Inhibitor were purchased from Invitrogen (Carlsbad, CA). The Phospho-Akt (Ser473) antibody was purchased from Cell Signaling Technology (Beverly, MA). The pT3.Cas9-UTRglobin plasmid was a gift from Prof. Yonghua Sun, Institute of Hydrobiology, Chinese Academy of Sciences.

### Zebrafish

Zebrafish were raised following standard zebrafish husbandry guidelines (Westerfield 2000). Embryos were obtained by natural cross and staged following Kimmel *et al*. (Kimmel, Ballard et al. 1995). All embryos were raised in the standard E3 embryo medium (Westerfield 2000) until 3 dpf. The control embryo medium (containing 0.2 mM [Ca^2+^]) and the induction embryo medium (containing 0.001 mM [Ca^2+^]) were prepared following a previously reported formula (Dai, Bai et al. 2014). In some experiments, 0.003% (w/v) N-phenylthiourea (PTU) was added to the media to inhibit pigmentation. The *Tg(igfbp5a:GFP)* fish line was generated in a previous study (Liu, Dai et al. 2017). All experiments were conducted in accordance with the guidelines approved by the University of Michigan Institutional Committee on the Use and Care of Animals.

### Fluorescence-activated cell sorting (FACS), RNA-seq, and quantitative real-time PCR (qPCR) analysis

Isolation and sorting of NaR cells by FACS were performed as previously described (Liu, Dai et al. 2017). For RNA-seq, RNA was isolated from FACS-sorted NaR cells using a RNeasy Micro Kit (Qiagen, Valencia, CA). The integrity of the RNA samples was confirmed using the Agilent Bioanalyzer 2100 with an RNA 6000 Pico Kit (Agilent, Santa Clara, CA). All samples had an RNA Integrity Number (RIN) of 8 or greater. cDNA libraries were prepared using a SMART-Seq v4 Ultra Low Input RNA Kit (Singulomics, Palo Alto, CA). Briefly, the first-strand cDNA was synthesized, full-length ds cDNA was amplified by LD-PCR. The amplified cDNA was purified using an Agencourt AMPure XP Kit (Beckman Coulter Life Sciences) and validated using the Agilent 2100 Bioanalyzer (Agilent). An Illumina Library was prepared. Single-end, 50bp sequencing was performed on the Illumina Hi-Seq 4000 platform.

The quality control of raw reads was tested using FastQC. The raw reads were trimmed using the Trimmomatic (Bolger, Lohse et al. 2014) and filtered. The preprocessed reads were mapped to the genome sequence of zebrafish GRCz10 (GCA_000002035.3) using TopHat and the aligned reads were assembled into transcripts using Cufflinks. The assembled transcripts were merged with the reference annotation (Danio_rerio.Zv9.68) using cuffmerge. The SVA package in R was used for removing batch effects. The htseq-count function was used to detect the number of counts in BAM files. Differential expression analysis was performed using wilcox.test in R package. Heatmap of hierarchical clustering was created using the pheatmap R package. The Clusterprofilter package in R was used to perform GO and pathway analysis. To validate the RNA-seq results, NaR cells were FACS sorted and RNA isolation was carried out. cDNA was synthesized using Superscript III Reverse Transcriptase. qPCR was performed on an ABI 7500 fast Real-Time PCR system (Applied Biosystems, Foster City, CA) using SYBR Green (Bio-Rad). PCR primers are listed in Supplementary Table S2.

### Phylogenetic analysis of the Fkbp protein family

Phylogenetic tree of the FKBP family in humans (hs), zebrafish (dr) and Drosophila (dm) was constructed using the Maximum Likelihood method based on the JTT matrix-based model (Jones, Taylor et al. 1992). The tree was constructed using MEGA7 (Kumar, Stecher et al. 2016).

### Drug treatment

Drugs used in this study were dissolved in DMSO and further diluted in water. Drug treatments began at 3dpf. The fish were subjected to immunostaining after one day treatment or NaR cell quantification after two days of treatment.

### NaR cell measurement

*Tg*(*igfbp5a*:*GFP*) larvae were imaged under a Leica MZ16F stereo microscope (Leica, Hamburg, Germany) equipped with a QICAM 12-bit Mono Fast 1394 Cooled camera (QImaging, Surry, BC, Canada). Lateral view images were taken and GFP-positive NaR cells were quantified as described (Liu et al., 2017).

### CRISPR/Cas9-mediated F0 gene deletion

Four sgRNAs were used to perform transient deletion of *fkbp5* and *fkbp7* following (Wu, Lam et al. 2018). The sgRNAs were synthesized by *in vitro* transcription based on a published method (Shao, Guan et al. 2014). Cas9 mRNA was synthesized by *in vitro* transcription using the pT3.Cas9-UTRglobin plasmid as template. sgRNAs (40 ng/μl) were mixed with Cas9 mRNA (400 ng/μl) and co-injected into *Tg(igfbp5a:GFP)* embryos at one cell stage. The injected embryos were raised in E3 embryo medium. To induce NaR cell reactivation, fish were transferred to the control and induction embryo medium at 3 dpf as previously reported (Liu, Dai et al. 2017).

### Tol2 transposon-mediated transgenesis assay

A Tol2 transposon-mediated genetic mosaic assay was used to target expressed a transgene in a subset of NaR cells (Liu, Xin et al. 2018). *BAC(igfbp5a:myrAkt-mCherry)* DNA and Tol2 mRNA were mixed and injected into 1-cell stage *Tg(igfbp5a:GFP)* embryos. The embryos were raised and subjected to various treatments. Cells co-expressing mCherry and GFP were identified and the cell proliferation index was determined as previously reported (Liu, Xin et al. 2018).

### Whole-mount immunostaining

Zebrafish larvae were fixed in 4% paraformaldehyde, permeabilized in methanol and subjected to whole mount immunostaining of phosphor-Akt as described previously (Dai, Bai et al. 2014).

### Statistical analysis

Statistical analyses were performed in consultation with the University of Michigan’s Consulting for Statistics, Computing and Analytics Research (CSCAR) team. Cell proliferation index data were analyzed using pairwise Chi-square tests. Data are shown as mean ± standard deviation (SD). Statistical analyses between two groups were performed using unpaired two-tailed *t* test. When comparing the means of multiple groups to a control group, Dunnett test was used to correct for multiple comparisons after one-way ANOVA. Statistical significances from all tests were accepted at *p* < 0.05 or higher.

## Supplementary Figure

**Figure S1.**
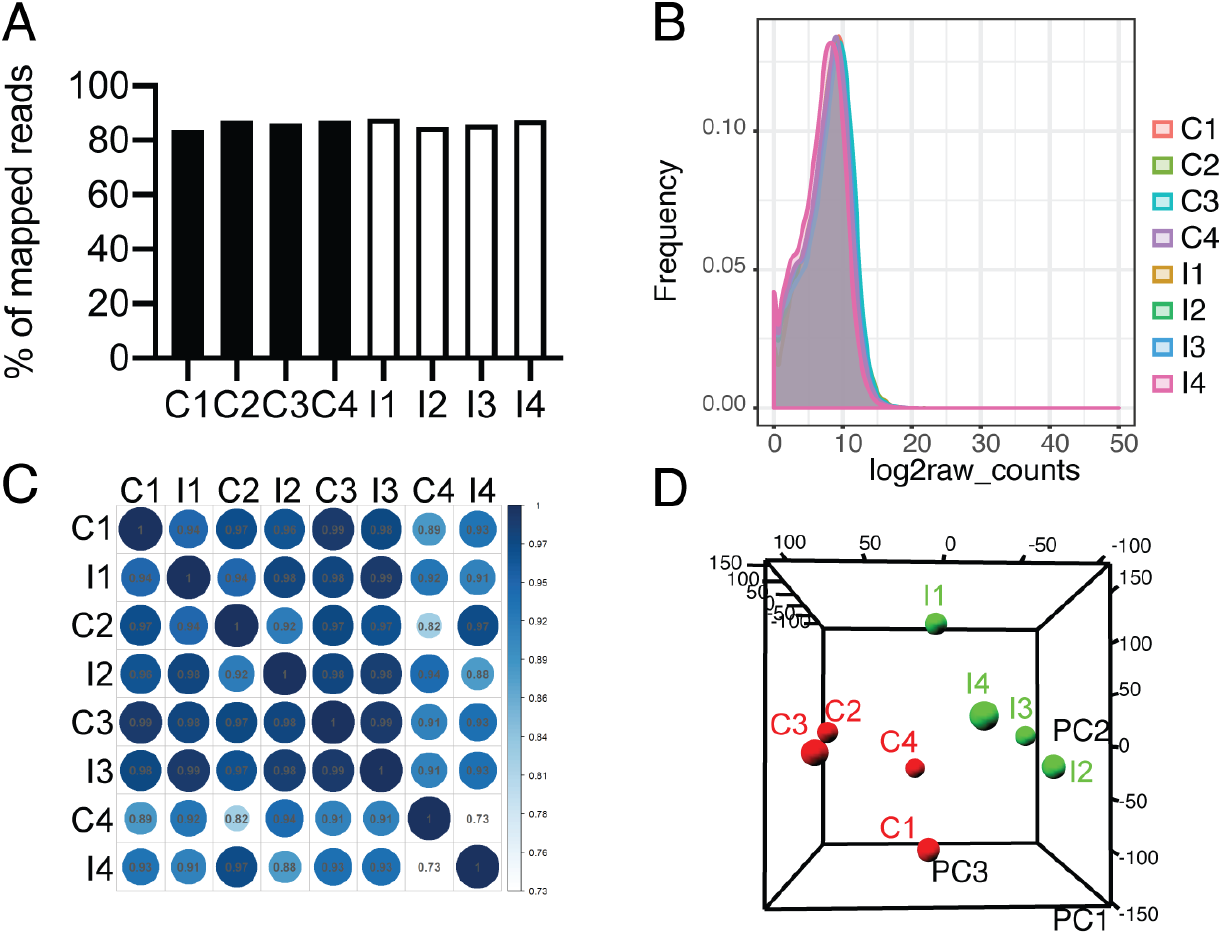
RNA-seq data quality. (A) Percentage of read mapping. (B) Distribution of count values for genes expressed in the four control (C) and four induction groups (I) of NaR cells. (C) Correlation coefficient analysis. Values show squared Pearson correlation (R^2^). (D) Principal component analysis (PCA) plot of the control (C) and induction groups of cells. Principal component 1 (PC1), and principal component 2 (PC2) were used for analysis.

**Figure S2.**
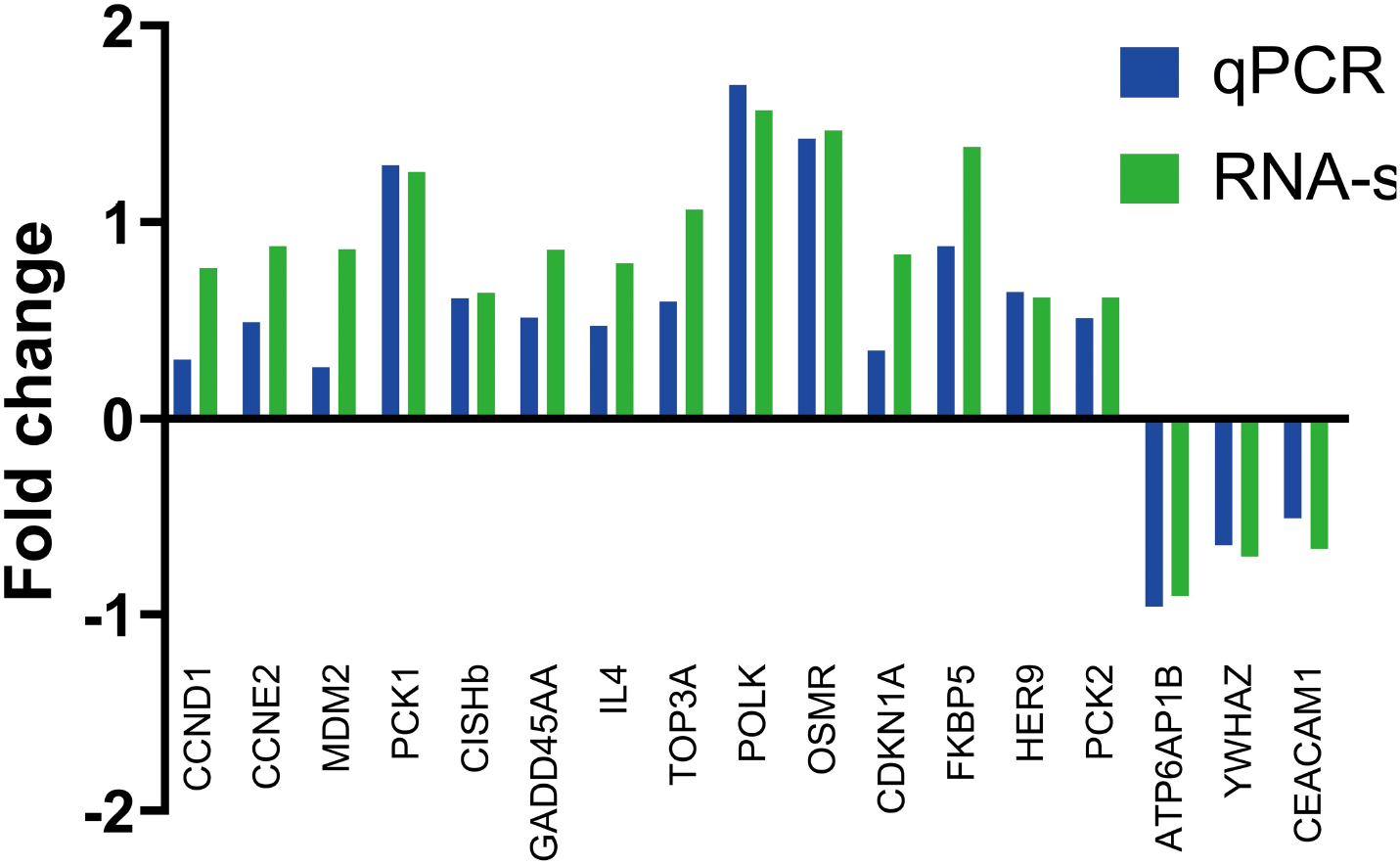
Changes in gene expression. NaR cells were isolated by FACS sorting as described in Fig. 1A. The mRNA levels of the indicated genes were measured by qPCR (blue) or RNA-seq (green) and shown as the ratio between the induction groups and the control groups.

**Figure S3.**
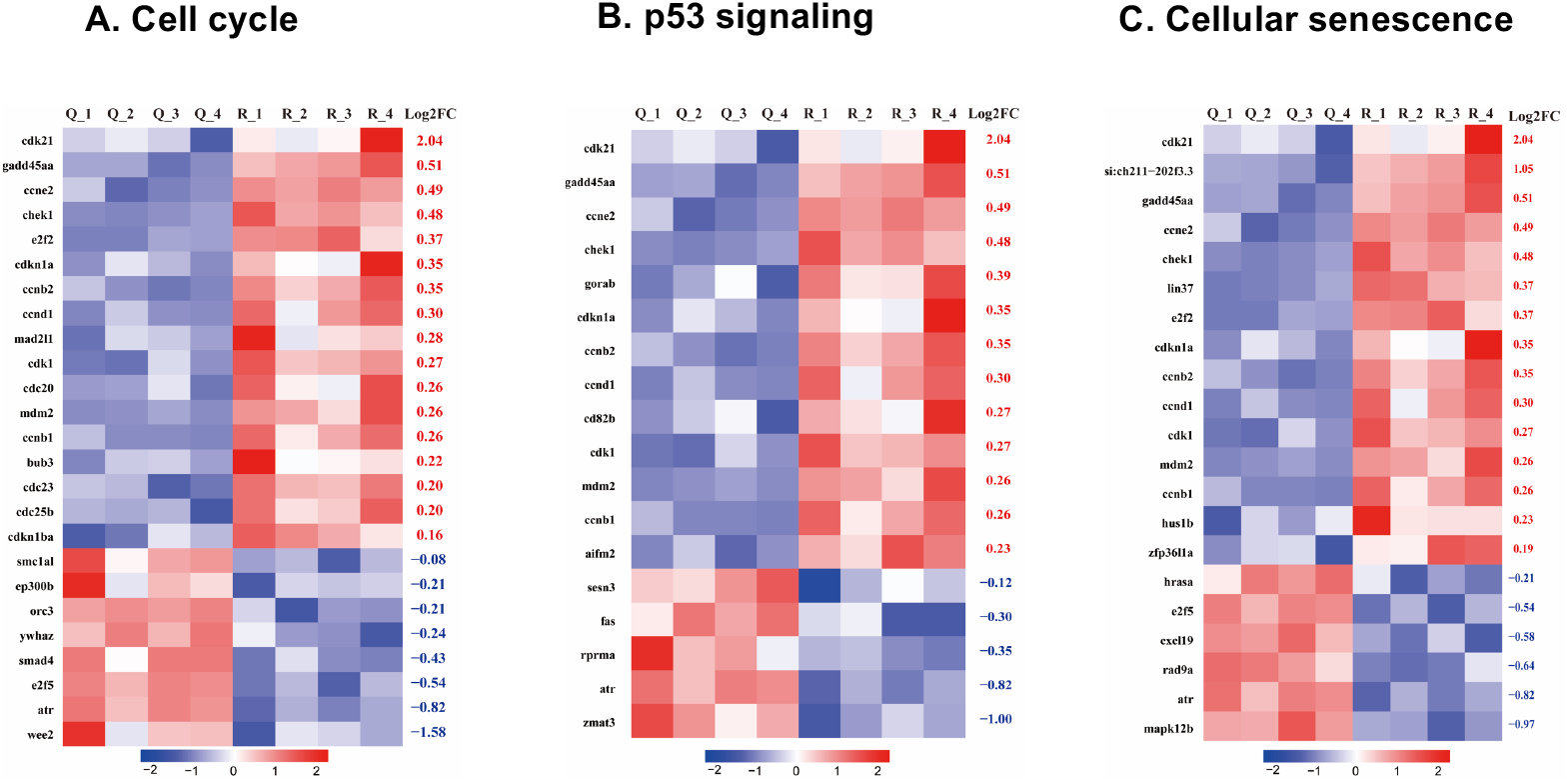
Heatmaps of enriched genes in cell cycle (A), p53 pathway (B), and cellular senescence (C) in the four control (C1-C4) and induction (R1-R4) groups of cells. Note the significant overlaps in these 3 categories.

**Figure S4.**
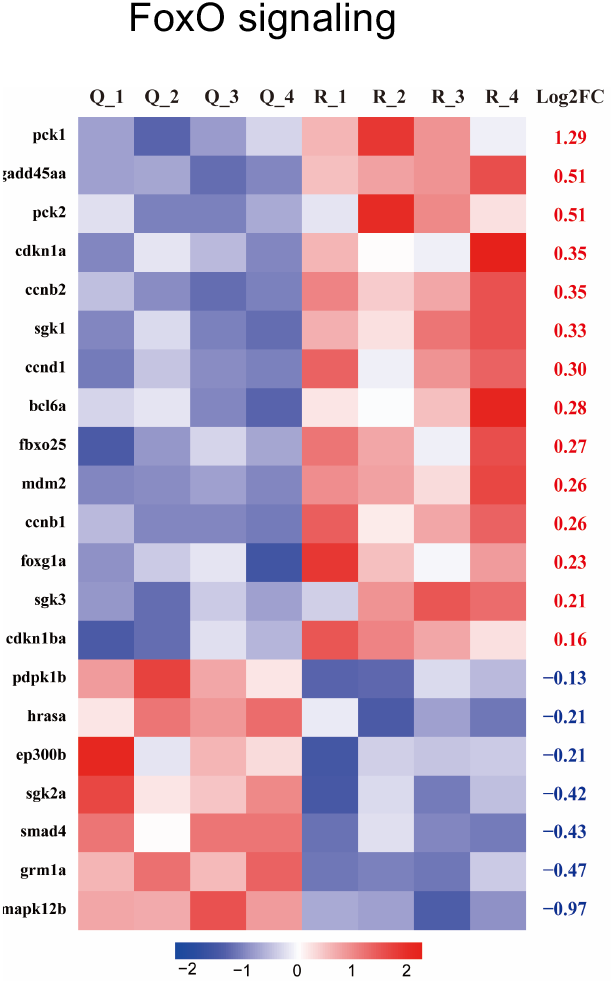
Heatmap of enriched genes in the Foxo signaling pathway in the four control (C1-C4) and four induction (R1-R4) groups of NaR cells.

**Figure S5.**
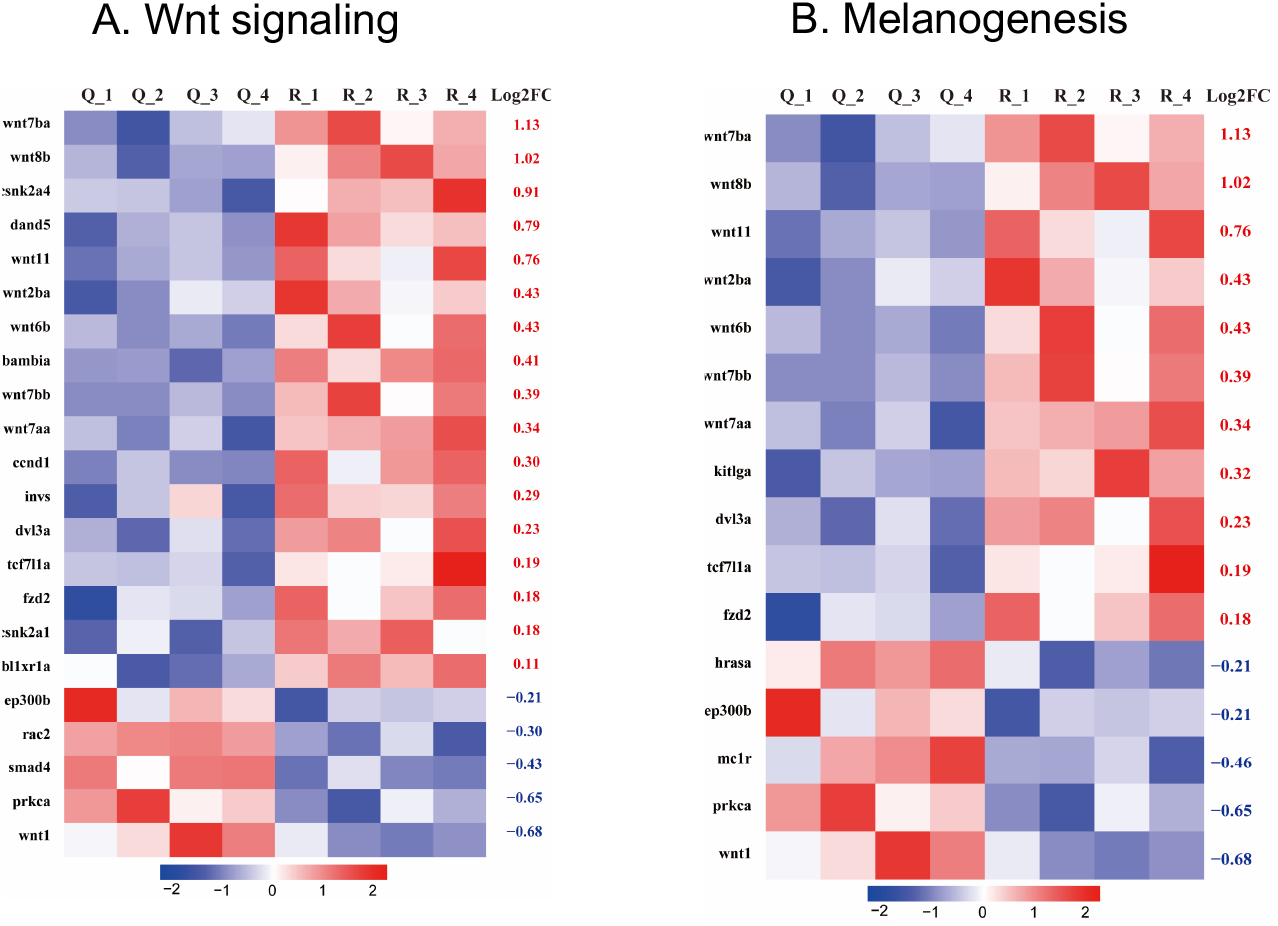
Heatmaps of enriched genes in Wnt signaling (A) and melanogenesis (B) pathway in the four control (C1-C4) and four induction (R1-R4) groups of NaR cells. Note the significant overlap in EDGs in these 2 categories.

**Figure S6.**
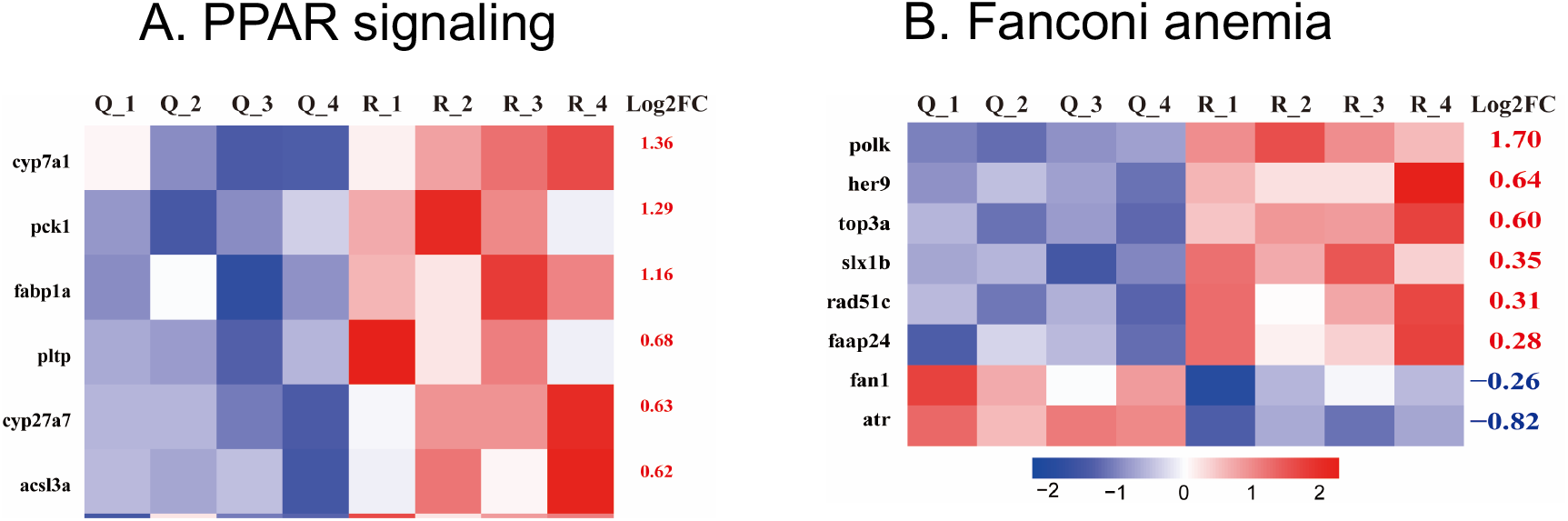
Heatmaps of enriched genes in the PAAR signaling (A) and Fanconi anemia pathway (B) in the four control (C1-C4) and four induction (R1-R4) groups of NaR cells.

