## Supplementary material for "FK506 binding protein 5 regulates cell quiescence-proliferation decision in zebrafish epithelium": Table S2

**Table S2. Genes and primers used in qPCR**

| Gene | Forward primer sequence (5’-3’) | Reverse primer sequences (5’-3’) |
| --- | --- | --- |
| *ccnd1* | CTGCGCAAACACGCCCAGAC | TACCGCTGCAGCAACACTGCC |
| *ccne2* | ACTGGACACTGCGGACAAAG | CTGCGATTTTTGTTGTGGTGC |
| *mdm2* | CCGAGGCAGACTACTGGAAG | CGAAGGTTGTGTTGGGAGTT |
| *pck1* | ATTTCAGCCGTGAACTGAACC | CGTAATGCAGATTAACGTGTGTG |
| *cishb* | TGCACCACTATTCATCCGCA | GTAAAGCAGCAGTGGTCGTG |
| *gadd45aa* | CACTGACGACGACGATGTGA | TTGATGTCGTTCTCGCAGCA |
| *il4* | ATTGGTCCCCGTTTCTGAGTC | TTCCTGCTTGGCAGAGAGTT |
| *top3a* | CCTGAACCTTACCCGTCTG | CCTGTAAACCGTTGGTGTATTTGG |
| *polk* | ACAAACCGATGGCAGTGGG | GACTCATAGGCATGAAATTAGGG |
| *osmr* | GTTTGGGACGAGAGAGCGAA | CCTGAACGGACTCCTGTACG |
| *cdkn1a* | ATGCAGCTCCAGACAGATGA | CGCAAACAGACCAACATCAC |
| *fkbp5* | GGGAAATGGACCTCAAAGAGA | CGATCCGCTGGTACTGAATTA |
| *her9* | TTCAGATGAGCGCAGCCTTG | TCCCTCGCAGGTAGACAGAA |
| *pck2* | AAGTTCACGTGGTCACAGGC | CCTCCGCTGGGGATAGGAAT |
| *atp6ap1b* | CCAGGCTGATCTAGCAAGCA | GTCCAGTTGCGAGCAGAAAC |
| *ywhaz* | GAGCGACCACTCTCACTCAC | CGAGCTCCCACCACATTCTT |
| *ceacam1* | GACATGACCCAGGCCTATAAC | CAGCTGAAGGTCACACTATCTC |
| *fkbp7* | ACATGTGTCCAGGCGAGAAG | TGCGTGGTCCTCTGCTTATG |
| *18s* | AATCGCATTTGCCATCACCG | TCACCACCCTCTCAACCTCA |
